# Floristic composition, phenology, and conservation value of four peat bogs in Bucovina, with the presence of *Betula nana*

**DOI:** 10.64898/2026.09.22.752958

**Authors:** Marian Trisciuc

## Abstract

This paper presents a comparative analysis of the floristic composition and site characteristics of four peat bogs in Bucovina, Romania: Poiana Stampei, Romanesti, Saru Dornei, and Gaina-Lucina. The research was based on phytosociological relevés on 25 m^2^ plots and direct field phenological observations, on six field visits from May to August 2026. Vegetation was characterised using the Braun–Blanquet method, and floristic similarity between sites was assessed with the Sørensen and Bray–Curtis indices.

All four plots shared a common core of taxa characteristic of peatland vegetation: *Sphagnum* spp., *Carex rostrata, Drosera rotundifolia, Eriophorum vaginatum*, and *Vaccinium* species. Species richness was 13 taxa at Poiana Stampei, Romanesti, and Saru Dornei, and 12 at Gaina-Lucina. Romanesti and Saru Dornei showed the highest floristic similarity (descriptive values, not statistically tested, given a single relevé per site), while Gaina-Lucina differed most markedly, not through species richness, which was similar across sites, but through species identity and through the presence of *Betula nana*, a glacial relict absent from the other sites. The results provide a descriptive basis for future research on the floristic composition and conservation of these habitats.

## 1. Introduction

Active raised bogs are peatland ecosystems characterised by the accumulation of organic matter under water-saturated conditions and by the development of vegetation adapted to the specific environmental conditions of these ecosystems (Frink et al., 2014). They develop under acidic, nutrient-poor conditions, with water input provided mainly by precipitation, characteristics that shape the structure and composition of the vegetation (Frink et al., 2014). Peat bog vegetation is associated in particular with the development of peat mosses, the genus *Sphagnum* playing an important role in the structure of these communities (Frink et al., 2014). In Romania, raised bogs and other peatland habitats are classified into distinct habitat types according to their ecological and floristic characteristics (Donita et al., 2005).

In the Eastern Carpathians, raised bogs are important components of the wet montane landscape, and the Dorna Depression and neighbouring areas in the northern Eastern Carpathians include several peatland areas with distinct floristic and ecological features (Pop, 1960; Frink et al., 2014). The vegetation of these ecosystems is influenced by substrate characteristics, hydrological regime, altitude, microrelief, and the structure of the woody vegetation, which can produce differences between peat bogs even when they belong to the same general habitat type.

The *Sphagnum* carpet represents one of the main structural components of peat bog vegetation, being associated with numerous herbaceous species, shrubs, and, locally, woody species. The importance of the genus *Sphagnum* for the structure and functioning of peat bogs is supported by research carried out in several Carpathian regions, including the Apuseni Mountains (Sass-Gyarmati et al., 2025).

Floristic composition may vary with local conditions, and differences between peat bog communities can reflect both site-specific features and the developmental stage of the vegetation (Frink et al., 2014; Robroek et al., 2017). The vegetation of montane peat bogs in the Romanian Carpathians has been analysed in earlier phytosociological studies, including for raised bogs in the Eastern Carpathians (Coldea & Plamada, 1989; Coldea et al., 2015; Stoica et al., 2022).

Active raised bogs are included among habitats of Community interest, and priority habitat 7110* is associated with active raised bogs within the Natura 2000 network. The conservation value of these ecosystems is determined both by their ecological characteristics and by their sensitivity to changes in the hydrological regime, to fragmentation of the habitat area, and to structural changes in the vegetation (Council Directive 92/43/EEC, 1992; Donita et al., 2005; Frink et al., 2014).

Several peat bogs with distinct floristic and site-related features exist in Bucovina (Bleahu et al., 2006). A number of these have been investigated over time, among which Poiana Stampei, Romanesti, and Saru Dornei were included in earlier phytosociological research on the oligotrophic vegetation of peat bogs in the Romanian Carpathians (Coldea & Plamada, 1989). For Poiana Stampei, dedicated studies have described the specific plant associations (Danu & Chifu, 2007) and the relationship between vegetation type and peat layer thickness (Gheorghe et al., 2006). Subsequent research on *Pinus sylvestris* vegetation in Eastern Carpathian peat bogs has also provided a regional comparison framework for these communities (Stoica et al., 2022). This information offers a reference point for interpreting the present floristic composition, although differences between historical and present-day data cannot automatically be interpreted as changes across the entire peat bog, since the plots and sampling methods are not identical.

A particularly important element of the peat bog flora of Bucovina is *Betula nana*. The species is regarded as a glacial relict in the flora of Romania, known with certainty from only two peat bogs in the Eastern Carpathians: Gaina-Lucina and Tinovul Luci (Pop, 1960; Stefan & Oprea, 2001; Borbély & Indreica, 2019). For Gaina-Lucina, previously published data have highlighted the conservation importance of the site and the presence of floristic elements of particular value, including *Betula nana* (Ursu et al., 2017).

The aim of the present study was to compare the floristic composition, taxon cover-abundance, and observed phenological succession in four peat bogs in Bucovina, based on data obtained directly in the field in 2026. The four peat bogs, Poiana Stampei, Romanesti, Saru Dornei, and Gaina-Lucina, were selected to combine sites with a documented phytosociological history (the first three) with the site of highest conservation value in the region, owing to the unique presence of *Betula nana* (Gaina-Lucina), thereby providing both a reference point for comparison with previously published data and a case of high conservation interest. The site characteristics of the plots investigated and the floristic similarity among them were also examined.

Three of the four peat bogs investigated, namely Poiana Stampei, Romanesti, and Saru Dornei, have previously been the subject of phytosociological research. The comparison of present-day data with previously published information is used in this study for orientation purposes only, given the differences in methodology, relevé location, and taxonomic resolution.

The research was purely observational and non-destructive in nature: taxon identification, cover-abundance estimation, and phenological observations were carried out directly in the field, without collecting plant material, and photographs taken during fieldwork were used as auxiliary material to verify the observations.

## 2. Materials and methods

### 2.1 Study area

The study was carried out in four peat bogs in the northern Eastern Carpathians, in the Bucovina region: Poiana Stampei, Romanesti, Saru Dornei, and Gaina-Lucina peat bogs. The four plots differ in site conditions, reflected mainly in altitude, soil type, local slope, and other site characteristics, allowing a descriptive comparison of floristic composition and vegetation structure. The location of the four plots investigated is shown in Figure 1.

**Figure 1.**
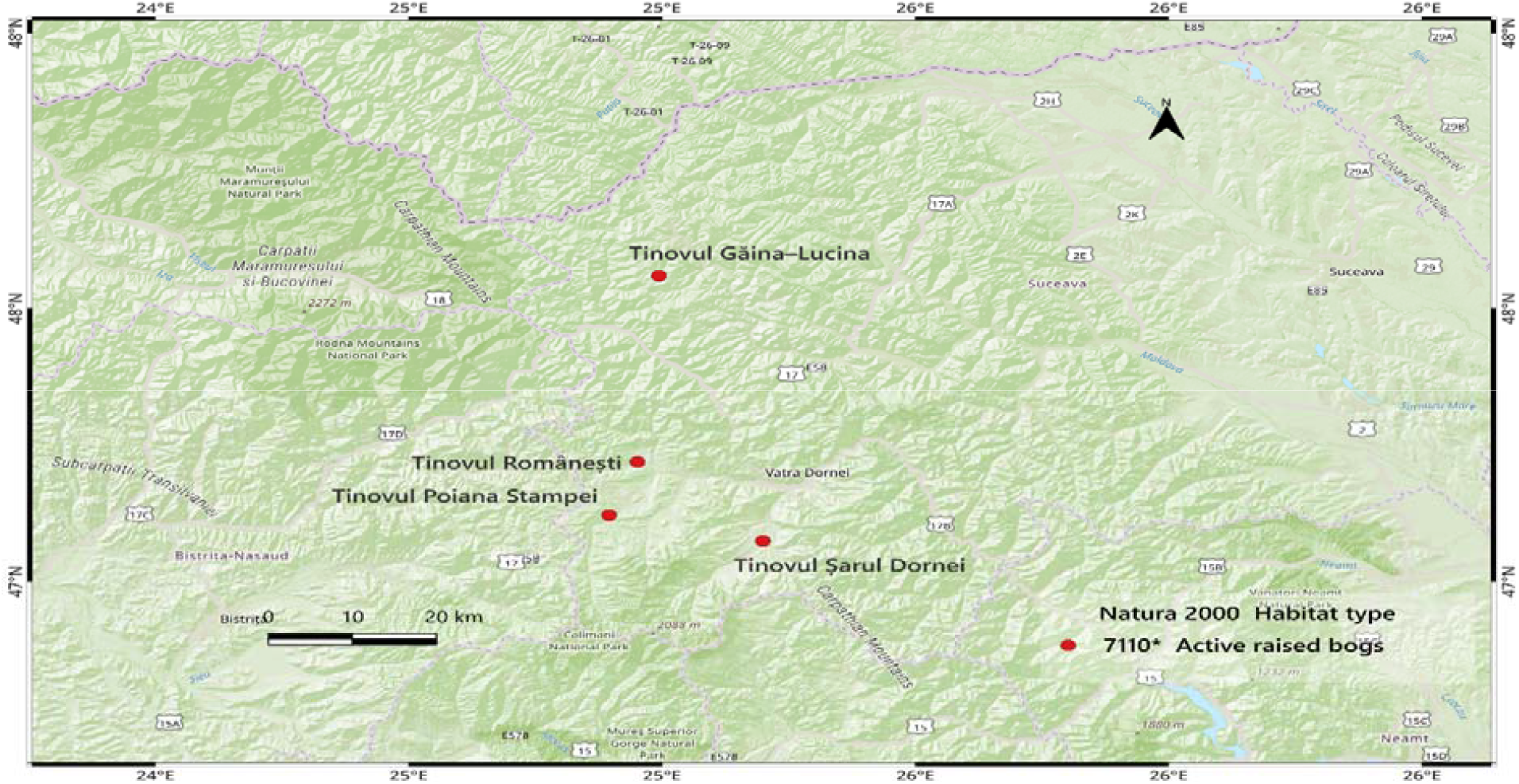
Location of the four peat bogs investigated in Suceava County (map produced in QGIS by the author).

For each plot, geographic coordinates, altitude, topographic position, slope, soil type, and relevé area were recorded. Information on protection status and conservation documentation for the sites was consulted as contextual sources for interpreting the results.

Floristic observations were carried out in 2026. A single 25 m^2^ plot was analysed in each peat bog, placed in an area considered representative of the peat bog vegetation. The choice of plot size aimed to obtain floristic data comparable across sites while, at the same time, limiting the impact of the research on the *Sphagnum* carpet and associated vegetation.

Repeating relevés over several plots would have required additional field visits and a more extensive disturbance of the vegetation, which is why the sampling effort was limited to one standardised plot per peat bog. The site characteristics of the four plots, including moss layer cover, are summarised in Table 1.

**Table 1.** Site characteristics of the four plots investigated.

| Characteristic | Poiana Stampei | Romanesti | Saru Dornei | Gaina-Lucina |
| --- | --- | --- | --- | --- |
| Coordinates (lat., long.) | 47.2997° N, 25.1136° E | 47.3754° N, 25.1686° E | 47.2586° N, 25.3548° E | 47.6477° N, 25.1958° E |
| Altitude (m) | 920 | 860 | 900 | 1165 |
| Soil type | Histosol | Gleysol (peaty subtype) | Gleysol (peaty subtype) | Histosol |
| Topographic position | flat | flat | flat | flat |
| Slope (°) | 4 | 3 | 3 | 5 |
| Relevé area (m<sup>2</sup>) | 25 | 25 | 25 | 25 |
| Canopy closure | 0.5 | 0.4 | 0.3 | 0.5 |
| Moss layer cover (%) | 75 | 80 | 90 | 70 |

Note. The soil type indicated in the management documentation available for the sites investigated (Ministry of Environment, Waters and Forests [MMAF], 2016a, 2016b, 2016c, 2016d) was cross-checked against field observations, based on the morphological characteristics of the soil (presence and thickness of the peat horizon), for the area of each relevé. At the whole-site level, the management plans indicate, for Poiana Stampei, Romanesti, and Saru Dornei, a predominance of hydromorphic soils, alongside Histosols and Cambisols; the distinction between Histosol and Gleysol (peaty subtype) was made on the basis of field morphological characters, without laboratory analyses that could further confirm the classification. Canopy closure was estimated visually in the field, on a scale from 0.1 to 1.0 (where 1.0 corresponds to a fully closed canopy).

Gaina-Lucina is the plot located at the highest altitude, 1165 m, the difference relative to the other plots being approximately 245–305 m.

### 2.2. Phytosociological relevé and cover estimation

Floristic composition was analysed through one 25 m^2^ phytosociological relevé in each of the four peat bogs, a plot size at the upper end of the 16–25 m^2^ range recommended for sample plots in mire and peat bog habitats (Frink et al., 2014). This size was also chosen so as to remain within the boundaries of the active-vegetation patches of habitat 7110*, which occupy small areas within the four peat bogs investigated; a larger relevé would have risked exceeding the boundaries of the target habitat. Taxon cover-abundance was estimated using the Braun–Blanquet method (Braun-Blanquet, 1964; Mueller-Dombois & Ellenberg, 1974).

Identified taxa were recorded in the floristic table together with the corresponding Braun–Blanquet values. For *Sphagnum*, identification was not extended to species level for all specimens observed. For this reason, the taxon was recorded as *Sphagnum* spp., avoiding the assignment of specific determinations that had not been sufficiently verified. For the same reason, a *Carex* specimen identified at Gaina-Lucina could not be determined with certainty to species level based on the characters observed in the field and the available photographic material, and was therefore recorded as *Carex* spp., in order to avoid reporting an uncertain identification.

For the moss layer, its estimated cover was recorded separately. This value expresses the proportion of the plot surface occupied by the moss layer and does not represent a transformation of the Braun–Blanquet values. For *Sphagnum* spp., the high cover observed in the field was therefore kept separate from the cover-abundance value.

For *Pinus sylvestris*, specimens showing morphological characters associated with the *turfosa* form were recorded in the field observations. For the comparative analysis of floristic composition, these were treated at the level of the species *Pinus sylvestris*.

### 2.3. Phenological observations and data processing

Phenological observations were carried out during six field visits, between 08 May 2026 and 23 August 2026. For each taxon, the phenological stage observed directly in the field was recorded, using the following categories: vegetative (Veg), budding (Bud), anthesis (Ant), fruiting (Fru), dispersal (Dis), and yellowing/senescence (Sen). The stage recorded represents the phenophase observed at the time of the visit and does not exclude the occurrence of a phenophase between two field visits.

Phenological observations were used to describe the succession of phenophases during the 2026 season. Because the systematic observations come from a single season, they are not used to establish multi-year phenological models.

The observations available from 2023 at Gaina-Lucina were used only for comparison, to highlight possible inter-annual differences relative to the systematic 2026 observations, without being integrated into the quantitative analysis of the 2026 relevés.

### 2.4. Floristic similarity analysis

Floristic similarity among the four plots was assessed using the Sørensen index (Sørensen, 1948), calculated from the presence and absence of taxa:

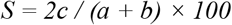

where a is the number of taxa in the first plot, b the number of taxa in the second plot, and c the number of taxa common to both.

To assess similarity based on cover-abundance, the Bray–Curtis index was used (Bray & Curtis, 1957). Braun–Blanquet values were transformed into an ordinal numerical scale following the scheme used by van der Maarel (1979): r = 1, + = 2, 1 = 3, 2 = 4, 3 = 5, 4 = 6, and 5 = 7. The same transformed values were used to calculate the Shannon–Wiener (H’) and Simpson (1−D) diversity indices for each plot, as indicative measures of diversity that combine taxonomic richness with cover-abundance structure; these values should be interpreted with caution, since the Braun–Blanquet scale is ordinal and the numerical transformation represents an approximation of actual abundance, not a direct quantitative measurement.

The Bray–Curtis index was calculated for each pair of plots based on the transformed values. Because each peat bog is represented by a single relevé, the indices are used to describe similarity among the plots investigated and not to statistically test differences between populations or sites.

## 3. Results

### 3.1. Site characteristics and floristic composition

The four plots analysed had the same 25 m^2^ area and were placed on terrain with gentle slopes, ranging between 3° and 5°, indicating a relatively uniform relief across the plots investigated. The main site-related differences were in altitude, ranging from 860 m at Romanesti to 1165 m at Gaina-Lucina. Poiana Stampei was located at 920 m, and Saru Dornei at 900 m. Regarding the soil substrate, the four plots were characterised by two soil types: Histosol at Poiana Stampei and Gaina-Lucina, and Gleysol (peaty subtype) at Romanesti and Saru Dornei.

The classification was based on morphological characteristics observed in the field and on information available in the management documentation for the areas investigated, in the absence of laboratory pedological analyses. Differences in floristic composition among the plots analysed were highlighted through the presence, absence, and cover-abundance values of the taxa identified. General aspects of the vegetation at the four sites are shown in Figure 2, and the cover-abundance values recorded for the identified taxa are presented in Table 2.

**Table 2.** Braun-Blanquet cover-abundance values for the taxa recorded at the four sites.

| Taxon | Poiana Stampei | Romanesti | Saru Dornei | Gaina-Lucina |
| --- | --- | --- | --- | --- |
| *Andromeda polifolia* | 1 | + | 1 | – |
| *Betula nana* | – | – | – | 2 |
| *Betula pubescens* | 1 | 1 | 1 | – |
| *Carex rostrata* | 1 | 1 | 1 | 1 |
| *Carex* spp. | – | – | – | 1 |
| *Drosera rotundifolia* | 1 | + | + | r |
| *Eriophorum angustifolium* | + | r | + | r |
| *Eriophorum vaginatum* | 1 | 1 | 1 | 1 |
| *Hylocomium splendens* | – | + | + | – |
| *Picea abies* | r | r | r | 1 |
| *Pinus sylvestris* | 3 | 3 | 3 | 3 |
| *Polytrichum* spp. | + | – | – | – |
| *Sphagnum* spp. | 4 | 4 | 5 | 4 |
| *Vaccinium oxycoccos* | 2 | 2 | 2 | – |
| *Vaccinium microcarpum* | – | – | – | 1 |
| *Vaccinium vitis-idaea* | + | + | + | 1 |
| *Vaccinium myrtillus* | 2 | 2 | 2 | 1 |
| Total taxa identified | 13 | 13 | 13 | 12 |
| Shannon–Wiener index (H') | 2.481 | 2.445 | 2.458 | 2.390 |
| Simpson index (1–D) | 0.910 | 0.904 | 0.905 | 0.902 |

**Table 3.** Inter-annual comparison (2023–2026) for taxa with recorded differences at Gaina-Lucina.

| Taxon | 2023 | 2026 |
| --- | --- | --- |
| *Vaccinium myrtillus* | anthesis observed | anthesis not observed |
| *Carex* spp. | not observed | present (BB = 1, vegetative stage) |

**Figure 2.**
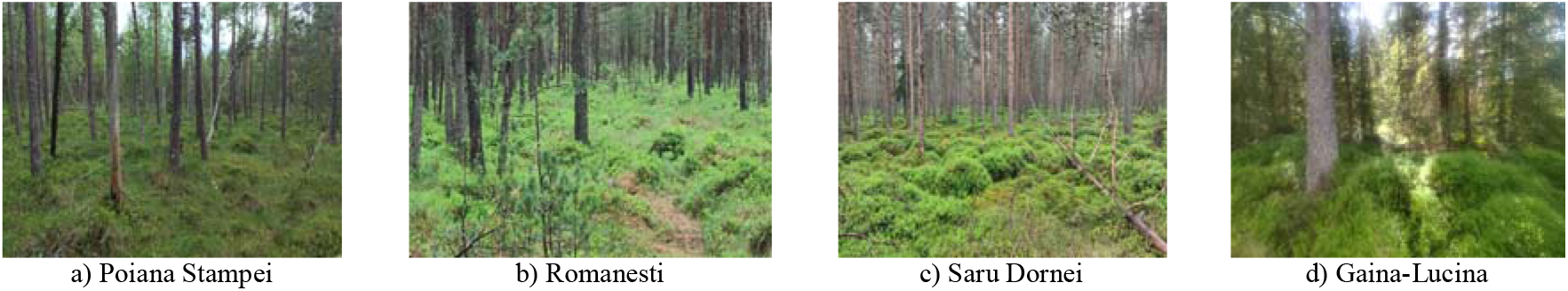
General aspects of the vegetation in the four peat bogs investigated. [Photo: Marian Trisciuc].

A total of 17 taxa were identified across the four relevés. Poiana Stampei, Romanesti, and Saru Dornei each had 13 taxa, and Gaina-Lucina had 12 taxa.

An important group of taxa was common to all four plots, namely Carex rostrata, Drosera rotundifolia, Eriophorum vaginatum, Eriophorum angustifolium, Picea abies, Pinus sylvestris, Sphagnum spp., Vaccinium vitis-idaea, and Vaccinium myrtillus. Vaccinium oxycoccos was recorded at Poiana Stampei, Romanesti, and Saru Dornei, while Vaccinium microcarpum was identified at Gaina-Lucina. Pinus sylvestris, including specimens with morphological characters associated with the turfosa form, was recorded at all four plots. Gaina-Lucina differed through the presence of Betula nana and Carex spp., taxa that were not recorded in the other three relevés. At the same time, Betula pubescens was identified at Poiana Stampei, Romanesti, and Saru Dornei. Several of the floristic elements observed are shown in Figure 3.

**Figure 3.**
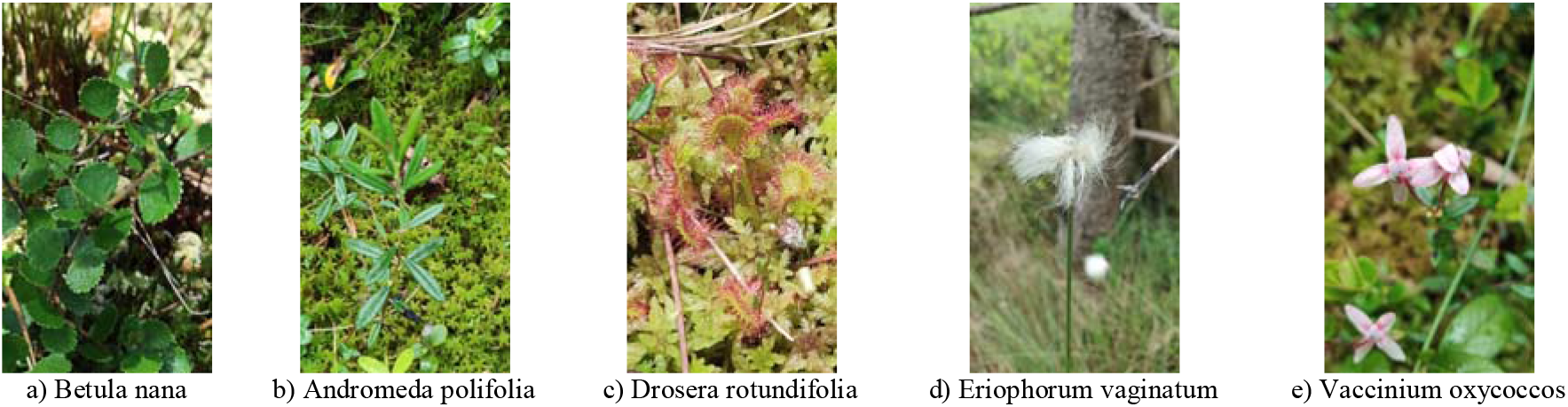
Characteristic taxa and vegetation elements observed in the field. [Photo: Marian Trisciuc].

Braun–Blanquet values highlighted the predominance of the *Sphagnum* layer. *Sphagnum* spp. had a value of 4 at Poiana Stampei and Romanesti, 5 at Saru Dornei, and 4 at Gaina-Lucina. *Pinus sylvestris* had a value of 3 at all four plots.

*Vaccinium myrtillus* had a value of 2 at Poiana Stampei, Romanesti, and Saru Dornei, and a value of 1 at Gaina-Lucina. *Vaccinium oxycoccos* had a value of 2 at the three plots where it was identified, while *Vaccinium microcarpum* had a value of 1 at Gaina-Lucina.

*Carex rostrata* and *Eriophorum vaginatum* had a value of 1 at all four relevés. *Eriophorum angustifolium* showed values ranging between r and +, while *Drosera rotundifolia* showed values ranging between r and 1. *Picea abies* had a value of r at Poiana Stampei, Romanesti, and Saru Dornei, and a value of 1 at Gaina-Lucina.

Moss layer cover was estimated at 75% at Poiana Stampei, 80% at Romanesti, 90% at Saru Dornei, and 70% at Gaina-Lucina. Locally, the *Sphagnum* carpet was observed to be continuous or nearly continuous. Canopy closure ranged from 0.3 at Saru Dornei to 0.5 at Poiana Stampei and Gaina-Lucina, and the observed values descriptively suggest a possible inverse association with moss layer cover. This observation nonetheless remains an unconfirmed hypothesis, which requires verification across a larger number of plots and through direct measurements of light conditions and other site factors. Thus, Saru Dornei, with the lowest canopy closure (0.3), showed the highest moss layer cover (90%), while Gaina-Lucina, with one of the highest canopy-closure values (0.5, equal to Poiana Stampei), recorded the lowest moss cover (70%). This observation must nevertheless be interpreted with caution, given the small number of plots investigated and the descriptive nature of the analysis.

Shannon–Wiener index values (2.390–2.481) and Simpson index values (0.902–0.910) were similar among the four plots investigated, indicating small differences in diversity calculated from the transformed cover-abundance values. Gaina-Lucina showed the lowest taxonomic richness (12 taxa) and the lowest values of both indices, although the differences relative to the other plots were small. The highest Shannon–Wiener value was recorded at Poiana Stampei (2.481), which had 13 taxa, a value comparable to those of the Romanesti and Saru Dornei plots. These results indicate a relatively similar diversity among the plots investigated, while the floristic particularity of Gaina-Lucina is mainly related to the identity of the taxa present and to floristic elements of conservation interest, aspects discussed in section 4.

### 3.2. Floristic similarity

The Sørensen index showed the highest similarity between Romanesti and Saru Dornei, with a value of 100%, the two plots having the same taxonomic composition in the dataset analysed.

Between Poiana Stampei and Romanesti, and between Poiana Stampei and Saru Dornei, similarity was 92.3%. Gaina-Lucina showed values of 72.0% relative to each of the other three plots (Figure 4). The Sørensen index reflects only the presence and absence of taxa and does not take into account differences in cover-abundance values.

**Figure 4.**
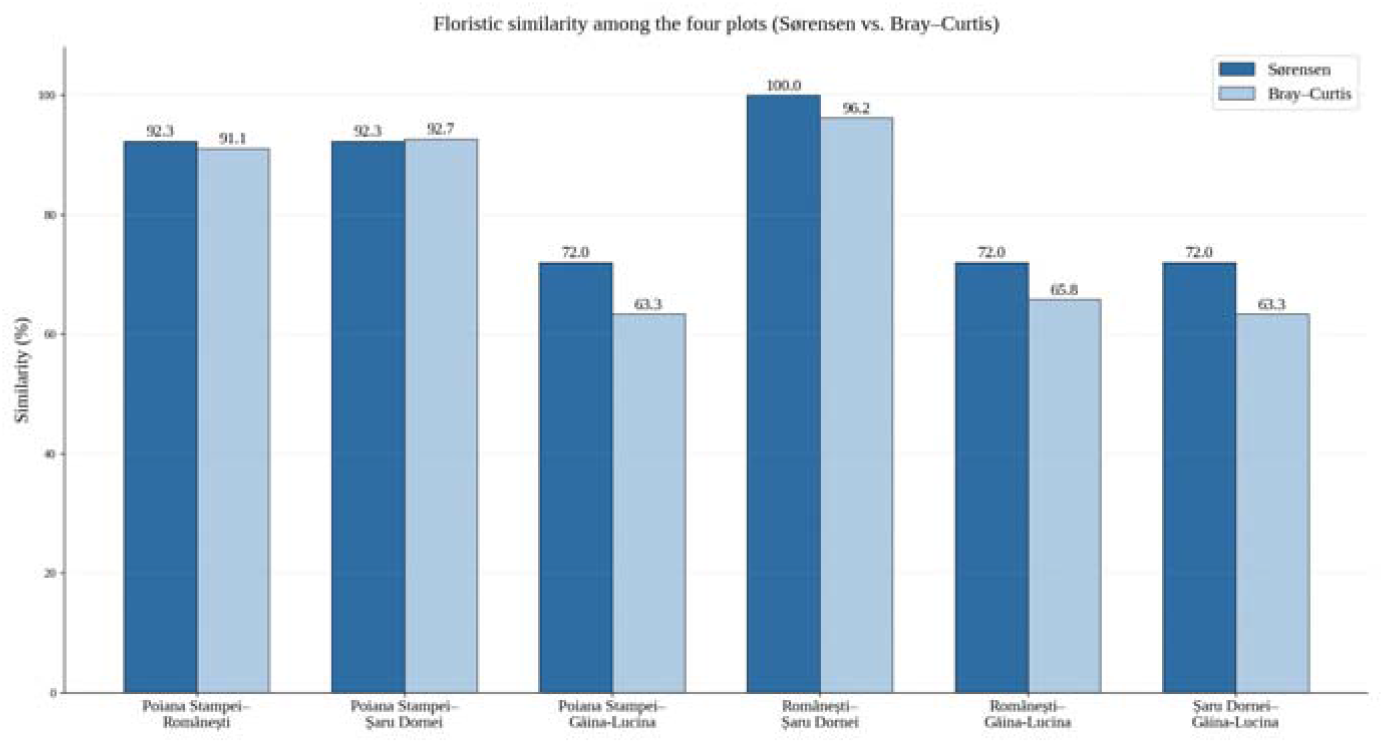
Floristic similarity (Sørensen and Bray–Curtis indices) among the four plots.

Based on the transformed Braun–Blanquet values, the Bray–Curtis index likewise showed the highest similarity between Romanesti and Saru Dornei, at 96.2%. Between Poiana Stampei and Saru Dornei the value was 92.7%, and between Poiana Stampei and Romanesti it was 91.1%. Gaina-Lucina showed lower values, between 63.3% and 65.8%. The results are shown in Figure 4.

Both similarity assessment methods showed the same general pattern: Romanesti and Saru Dornei had the highest floristic similarity among the plots investigated, while Gaina-Lucina differed most markedly from the other plots.

### 3.3. Observed phenological succession

Observations made from May to August 2026 revealed successions of phenological stages for most of the taxa monitored, including the absence of some taxa at the first visit, on 8 May, when the vegetation was still at the beginning of the season.

For *Andromeda polifolia*, at Poiana Stampei, Romanesti, and Saru Dornei, the taxon was not identified at the first visit (8 May), and the succession observed afterwards was Veg–Ant–Ant–Dis–Dis. For *Carex rostrata*, Poiana Stampei and Romanesti showed the succession Veg–Veg–Ant–Ant–Dis–Dis, Saru Dornei showed the succession Veg–Ant–Ant–Ant–Dis–Dis, and the same succession, Veg–Ant–Ant–Ant–Dis–Dis, was observed at Gaina-Lucina.

*Eriophorum vaginatum* and *Eriophorum angustifolium* were recorded in the vegetative stage at the first visit (8 May) and subsequently showed, at all four plots, successions from anthesis to dispersal and then to yellowing/senescence. *Drosera rotundifolia* was not identified at any plot at the 8 May visit, the taxon becoming visible only at the following visit; afterwards, the plant progressed from the vegetative stage to anthesis and then to yellowing over the course of the observation period. *Vaccinium oxycoccos*, and *Vaccinium microcarpum* at Gaina-Lucina, as well as *Vaccinium vitis-idaea*, showed successions of the type Veg–Ant–Ant–Fru–Fru–Fru, with the vegetative stage beginning at the 8 May visit.

*Vaccinium myrtillus* showed a different pattern between the first three plots and Gaina-Lucina. At Poiana Stampei, Romanesti, and Saru Dornei, budding was already observed at the first visit (8 May), followed by anthesis at the 14 June visit, with the succession Bud–Ant–Fru–Fru–Fru–Fru. At Gaina-Lucina, in 2026, no anthesis phenophases were observed, and the taxon was recorded in the vegetative stage at all six observation dates, including the 8 May visit.

*Picea abies, Pinus sylvestris*, and *Sphagnum* spp. were recorded in the vegetative stage throughout the monitoring period. *Hylocomium splendens* was likewise recorded in the vegetative stage at Romanesti and Saru Dornei. The complete set of observations is shown in Figure 5.

**Figure 5.**
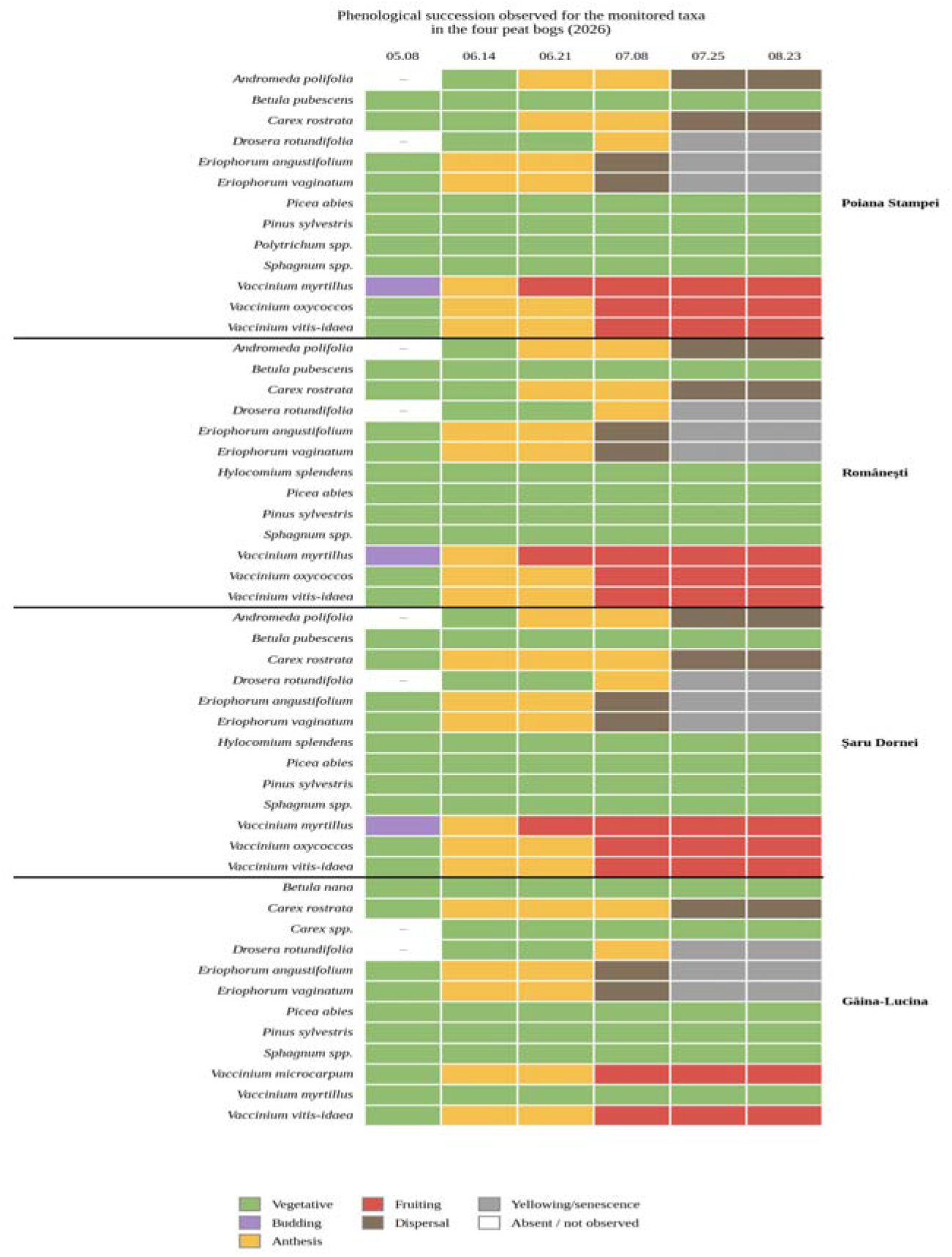
Phenological succession observed for the monitored taxa in the four peat bogs (2026).

The observations available from 2023 for Gaina-Lucina show two differences relative to the 2026 observations. In 2023, anthesis was observed in *Vaccinium myrtillus*, whereas in 2026 this phenophase was not observed during the monitoring period. Likewise, *Carex* spp. was not observed in 2023, whereas in the 2026 relevé it was recorded with a value of 1 and was observed in the vegetative stage throughout the monitoring period. These differences are consistent with inter-annual variability, but do not allow the identification of a temporal trend in the absence of continuous observations in the intervening years. The 2023 data are therefore used exclusively as a comparative and contextual element, without being interpreted as a time series and without allowing an assessment of any directional change in floristic composition between the two observation points.

### 3.4. Chorology and conservation status

For the taxa identified, sources on chorological element and rarity/conservation status used in this paper were consulted, namely Oltean et al. (1994) and Oprea (2005). Table 4 includes only taxa with a restricted chorological range (circumboreal, arctic-alpine, or Eurasian) or with a rarity/conservation status recorded in at least one of the sources consulted; taxa with a widespread range and no rarity status mentioned (for example *Betula pubescens, Carex rostrata*, or *Sphagnum* spp.) were not included. Information on *Betula nana* was supplemented with data from Dihoru & Negrean (2009) and Borbély & Indreica (2019). The summary is presented in Table 4.

**Table 4.**
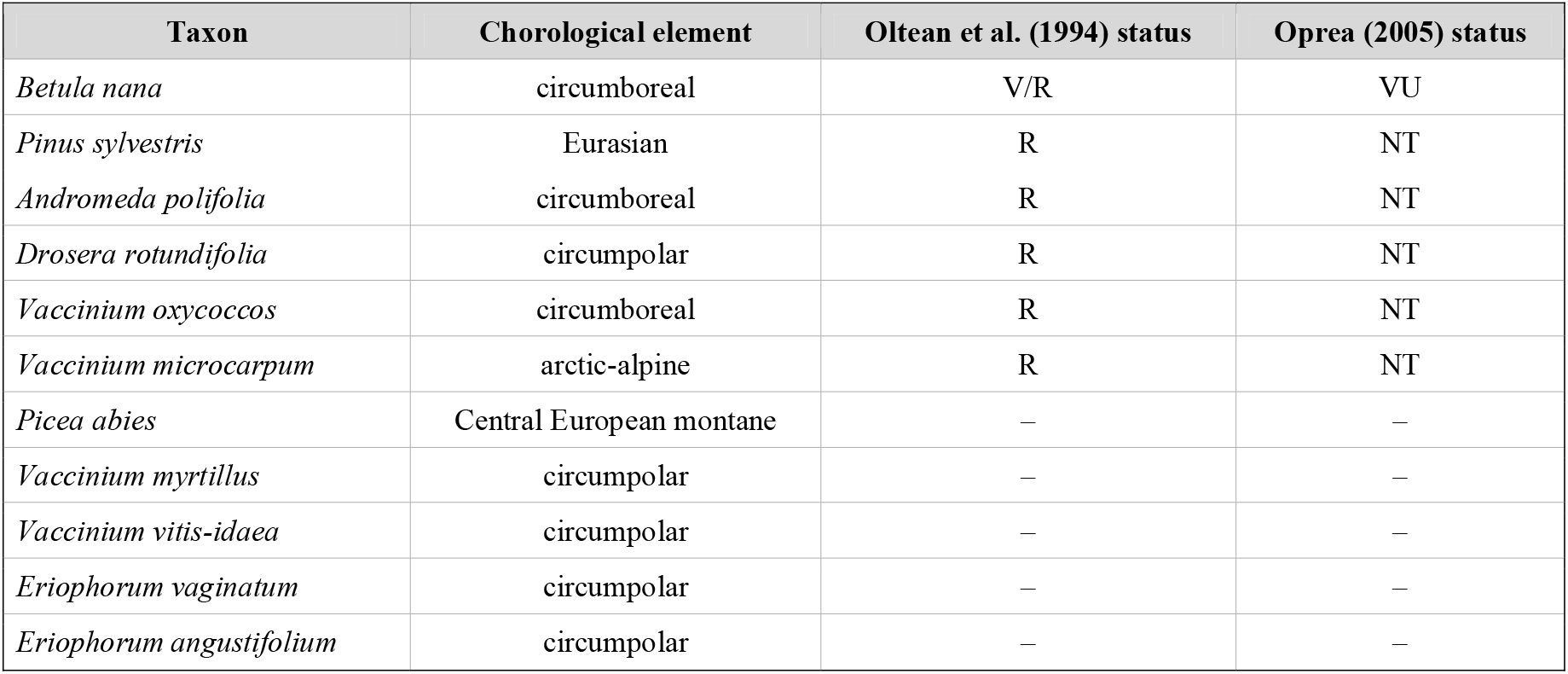
Chorological element and rarity/conservation status of the taxa identified (after Oltean et al., 1994 and Oprea, 2005).

*Betula nana* is the taxon of greatest conservation value among the floristic elements identified in this study. The species is regarded as a glacial relict, known with certainty from only two peat bogs of the Eastern Carpathians: Gaina-Lucina and Tinovul Luci (Pop, 1960; Borbély & Indreica, 2019). Dihoru & Negrean (2009) place it in the CR category at national level, based on IUCN criteria (IUCN, 2012), after it had previously been classified as V/R by Oltean et al. (1994), and as vulnerable (VU) by Oprea (2005). The remaining taxa with a circumboreal or arctic-alpine chorological element (*Andromeda polifolia, Drosera rotundifolia, Vaccinium oxycoccos, Vaccinium microcarpum*), as well as *Pinus sylvestris*, of Eurasian range, were classified as rare (R) by Oltean et al. (1994) and near threatened (NT) by Oprea (2005), without reaching a critical status.

The presence of *Betula nana* was confirmed in the plot analysed at Gaina-Lucina, where it was recorded with a value of 2 on the Braun–Blanquet scale. In the context of the present study, this presence constitutes a floristic element of conservation interest, although a single relevé does not allow the size or viability of the population across the entire peat bog to be assessed.

## 4. Discussion

The results obtained show the existence of a common floristic core among the four plots investigated. *Carex rostrata, Drosera rotundifolia, Eriophorum vaginatum, Eriophorum angustifolium, Sphagnum* spp., *Vaccinium vitis-idaea*, and *Vaccinium myrtillus* were recorded at all four relevés, consistent with the oligotrophic, acidic character of peat bog vegetation described in the specialist literature (Frink et al., 2014; Robroek et al., 2017). Within this common core, however, an ecological distinction can be noted: while *Sphagnum* spp., *Drosera rotundifolia, Eriophorum vaginatum*/*angustifolium*, and *Vaccinium* species of section *Oxycoccus* are typical peat bog specialists, *Vaccinium myrtillus* and *Hylocomium splendens* have a broader ecological amplitude, being common also in the understorey of acidic forests outside peat bogs; their presence indicates that the relevés included, alongside active peat bog vegetation, transitional microhabitats of the wooded-hummock type. Superimposed on this common core, however, are local particularities: although the four peat bogs belong to the same regional context, their taxonomic composition is not identical, highlighting the heterogeneous character of peat bog vegetation at the local scale. Differences among plots are not significant in terms of taxonomic richness (Poiana Stampei, Romanesti, and Saru Dornei each have 13 taxa, and Gaina-Lucina has 12), but rather in the identity of the taxa and their cover-abundance values. Gaina-Lucina shows the most pronounced particularity, through the presence of *Betula nana* and *Carex* spp., through the absence of taxa found at the other three plots, such as *Betula pubescens* and *Vaccinium oxycoccos*, and through the identification here of *Vaccinium microcarpum*. These differences must be interpreted at the level of the plots actually investigated, since each peat bog was represented by a single 25 m^2^ relevé, and the data characterise the floristic composition of the selected plots and do not allow the full floristic variability present in each peat bog to be estimated.

The very high similarity between Romanesti and Saru Dornei is supported both by the Sørensen index (100%) and by the Bray–Curtis index (96.2%) (section 3.2). The complete identity of the taxonomic list, reflected by the Sørensen index, nonetheless contrasts with the somewhat lower Bray–Curtis value, which indicates differences in the cover-abundance of the shared taxa, for example *Sphagnum* spp. had a value of 4 at Romanesti and 5 at Saru Dornei. The two methods thus provide complementary information: Sørensen highlights the similarity of taxonomic composition, while Bray–Curtis also captures quantitative differences. This high similarity must nonetheless be interpreted in relation to the point-based nature of the sampling, and the result cannot automatically be extended to the entire area of the two peat bogs.

Gaina-Lucina shows a more pronounced differentiation, reflected both in taxonomic composition and in cover-abundance values. This particularity can be discussed in relation to site-related differences: Gaina-Lucina is located at 1165 m altitude, compared with 860–920 m at the other three plots, and soil type, cross-checked with field observations based on morphological characteristics, differed among the plots investigated, Histosol at Poiana Stampei and Gaina-Lucina, and Gleysol (peaty subtype) at Romanesti and Saru Dornei. The correspondence observed between soil type and floristic similarity should be regarded as a working hypothesis, since the classification was not supported by laboratory pedological analyses, and the small number of plots investigated does not allow statistical testing of this relationship. It is notable that Poiana Stampei and Gaina-Lucina, both with Histosol, show the lowest floristic similarity among all pairs of sites (Sørensen 72.0%), whereas Romanesti and Saru Dornei, both with Gleysol, recorded the maximum floristic similarity (Sørensen 100%). This correspondence suggests a possible role of soil type in shaping the similarity between Romanesti and Saru Dornei, but, given the small number of sites and the absence of detailed pedological measurements, the relationship cannot be tested statistically and should be treated as a working hypothesis, not as a firm conclusion. The particularity of Gaina-Lucina should therefore be interpreted in the context of the overall site-related and ecological characteristics of the location, altitude, hydrological regime, microrelief, and vegetation structure being able to act simultaneously on the composition of peat bog communities (Frink et al., 2014; Robroek et al., 2017). The conservation importance of Eastern Carpathian peat bogs should also be viewed in the context of the regional distribution of these habitats: Mitof (2025) highlights the concentration of peatland habitats in the Eastern Carpathians and mentions the Gaina-Lucina site among the representative sites, information that provides additional regional context, without implying a direct causal relationship between habitat status and the floristic composition recorded in the relevé. In the absence of additional hydrological and pedological measurements (water level, pH, conductivity, moisture, peat layer thickness), field determination of soil type not having been extended to such parameters, the individual role of these factors cannot be separated, and the differentiation of Gaina-Lucina should be regarded as an association between the site characteristics observed and the floristic composition of the plot investigated, not as a demonstration of the effect of a single variable.

*Sphagnum* spp. is the dominant component of the moss layer at all four plots, with Braun–Blanquet values of 4–5 (section 3.1) and an estimated cover of 70–90%, confirming the importance of this component in the structure of the vegetation analysed. Locally, the carpet was observed to be continuous or nearly continuous, consistent with the specific character of peat bog vegetation (Frink et al., 2014; Robroek et al., 2017). Cover and cover-abundance values do not, however, allow extrapolation to the functional processes of the peat bogs: in the absence of additional measurements, the high presence of *Sphagnum* cannot be used to estimate productivity, peat accumulation rate, or hydrological dynamics. *Sphagnum* spp. should therefore be interpreted primarily as a dominant structural element of the plots analysed, without attributing to it quantitative ecological processes that were not measured directly. Data on canopy closure (section 3.1) suggest a possible role of shading on the development of the moss layer: the plot with the lowest canopy closure (Saru Dornei, 0.3) also showed the highest *Sphagnum* cover (90%), a pattern possibly consistent with the light requirements of some species within the genus *Sphagnum* (Bengtsson et al., 2021). The genus is, however, ecologically heterogeneous: species from typical hummock sections tolerate or prefer strong light, while others, also common in wooded peat bogs, are shade-tolerant, and the absence of species-level identification does not allow a uniform light response to be attributed. The relatively narrow range of variation in canopy closure (0.3–0.5) among the four plots also suggests moderate differences in illumination; hydrological regime and microtopography, not measured in this study, are factors that typically have a stronger influence on the development of the *Sphagnum* layer and could explain part of the pattern observed. The relationship therefore remains an unconfirmed hypothesis, which cannot be tested statistically based on only four plots.

The presence of *Pinus sylvestris* at all four plots, with a value of 3 on the Braun–Blanquet scale in each relevé, is consistent with previously published data on peat bog vegetation in the Eastern Carpathians, where the species is described as an important component of the *Vaccinio uliginosi-Pinetum sylvestris* and *Sphagno-Piceetum* associations (Coldea & Plamada, 1989; Chinan & Manzu, 2014; Stoica et al., 2022). Specimens with morphological characters associated with the *turfosa* form, anatomically documented in other Carpathian peat bog populations (Palla et al., 2021), were recorded in the field but treated at the level of the species *Pinus sylvestris*, since no anatomical or morphometric measurements were carried out that would allow a separate analysis. Agreement with the literature extends to other taxa as well: *Eriophorum vaginatum* and *Vaccinium vitis-idaea*, reported at high frequencies in previously studied *Pinus sylvestris* communities (Coldea & Plamada, 1989; Stoica et al., 2022), were recorded at all four plots in the present study, as was *Sphagnum* spp. Comparison with earlier studies nonetheless remains indicative, given differences in plot location, relevé size and number, methodology, and taxonomic resolution; a similar comparative approach was recently used for a montane peat bog community in the Bucegi Mountains (Mountford & Onete, 2024).

The absence of taxa mentioned in earlier floristic inventories should not automatically be interpreted as evidence of their disappearance from the peat bogs, since a single 25 m^2^ relevé cannot represent the entire area of a peat bog, and peat bog vegetation can show substantial spatial heterogeneity; differences between historical and present-day inventories may equally arise from the different location of the sample plots, from local vegetation structure, or from methodological differences. This limitation of the sampling design, namely a single 25 m^2^ plot per peat bog, was a deliberate choice: the plot size was set at the upper end of the 16–25 m^2^ range recommended for mires and peat bogs (Frink et al., 2014), aiming for data comparable across sites, keeping the relevé within the boundaries of the active-vegetation patches of habitat 7110*, which occupy small areas in the four peat bogs investigated, and, at the same time, reducing the impact of the research on the *Sphagnum* carpet and associated vegetation, a decision particularly relevant for sensitive habitats, where unjustified expansion of sampling can cause additional disturbance. The small number of plots therefore limits the extrapolation of the results to the level of the whole peat bog, but reflects a methodological decision aimed at minimising the impact on the habitat.

A further limitation is taxonomic: identification of *Sphagnum* to genus level was not extended to species level, although the recent literature offers both taxonomic checklists for European bryophytes (Hodgetts & Lockhart, 2020) and research dedicated to the diversity of the genus *Sphagnum* in Romania (Stefanut et al., 2026); these sources are used only as a reference for future research, not as evidence of the presence of particular species at the plots investigated. The same conservative approach was applied to the *Carex* specimen from Gaina-Lucina, recorded as *Carex* spp. because species-level determination could not be made with sufficient certainty, a speculative determination being best avoided, particularly in a study where the rarity status of a species could influence the interpretation of the results. Communities of epiphytic lichens, including *Usnea barbata*, were observed both on the trunk and on the branches of trees at all four plots investigated, but were not systematically inventoried or quantified, since they were not included in the initial sampling protocol; recording them could have added to the overall picture of the biodiversity of the plots, all the more so as *Usnea barbata* is considered a bioindicator of air quality and of forest habitat continuity, and this remains a direction for future research.

*Betula nana* remains the main floristic element of conservation interest identified at Gaina-Lucina. The species is regarded as a glacial relict in the flora of Romania, known with certainty from only two peat bogs of the Eastern Carpathians, Gaina-Lucina and Tinovul Luci (Pop, 1960; Borbély & Indreica, 2019), and its high conservation interest is reflected in successive classifications: V/R by Oltean et al. (1994), VU by Oprea (2005), and CR by Dihoru & Negrean (2009); these categories come from different sources and should not be confused with a new assessment carried out within the present study. The presence of the species, with a value of 2 on the Braun–Blanquet scale, confirms the floristic particularity of the plot, but a single relevé does not allow the size, age-class structure, or population trend to be estimated across the entire peat bog. Monitoring the *Betula nana* population at Gaina-Lucina, through repeated observations at the same plot and expanded inventorying in areas favourable to the species, therefore represents a priority direction for future research.

The 2026 phenological observations revealed a coherent seasonal succession for most of the taxa monitored, from anthesis to fruiting, dispersal, and, in some cases, yellowing or senescence, visible in particular in *Eriophorum, Carex, Drosera, Andromeda*, and *Vaccinium* (section 3.3). These successions must nonetheless be interpreted with caution: the six visits made between 08 May 2026 and 23 August 2026 allow observation of a single season, without supporting multi-year phenological models or attributing the differences observed to individual climatic or hydrological factors. The information available from 2023 at Gaina-Lucina provides an additional reference point: *Vaccinium myrtillus* was observed in anthesis in 2023, a phenophase not recorded in 2026, and *Carex* spp. was recorded only in 2026. These differences suggest possible inter-annual variability, but, coming from a single visit per peat bog in 2023 and in the absence of data for 2024–2025, they should be regarded as strictly descriptive, not as a demonstrated trend. Factors such as temperature, length of the growing season, altitude, and water availability may simultaneously influence phenology, but, in the absence of meteorological and hydrological data synchronised with the field observations, their individual role cannot be separated within the present study.

Overall, the four plots investigated show a common floristic core, over which relevant local differences are superimposed, the most evident particularity being that of Gaina-Lucina, expressed not through greater taxonomic richness, but through the identity of taxa of conservation relevance, in particular *Betula nana*. The results must be interpreted within the limits of the sampling design adopted, a single 25 m^2^ plot per peat bog, the point-based nature of the 2023 observations, phenological monitoring within a single season, identification of some taxa only to genus level, and the absence of direct hydrological and pedochemical measurements, limits that do not reduce the descriptive value of the observations but define the scope within which the results can be rigorously interpreted. The total area sampled (4 × 25 m^2^ = 100 m^2^) is also unrepresentative of the biodiversity of habitat 7110* at the regional scale; the results characterise exclusively the four points investigated and cannot be extrapolated to all peat bogs in Bucovina or in the Eastern Carpathians. The data obtained can thus provide a basis for further monitoring of the plots: repeating relevés, expanding sampling to additional plots, determining *Sphagnum* species, and integrating measurements of the hydrological regime and substrate characteristics would allow a more robust assessment of floristic variability and of the factors controlling vegetation structure, including the successional trajectory of these habitats (Kolari & Tahvanainen, 2023).

## 5. Conclusions

The research carried out at four peat bogs in Bucovina showed the existence of a common floristic core associated with peat bog vegetation, over which local differences among the plots investigated are superimposed. Although taxonomic richness was similar among the four relevés, differences in composition and cover-abundance revealed a more pronounced particularity at the Gaina-Lucina plot.

The high floristic similarity between Romanesti and Saru Dornei, shown both by the analysis of taxon presence and absence and by the transformed cover-abundance values, indicates a very similar floristic structure at the plots investigated. In contrast, the differentiation of Gaina-Lucina mainly reflects the identity of particular taxa and the local particularities of the plot analysed. The correspondence observed between soil type and floristic similarity represents a working hypothesis, which cannot be verified within the present study owing to the small number of plots and the absence of detailed pedological analyses.

*Sphagnum* spp. was the dominant structural component of the moss layer at all four plots investigated, with high cover-abundance and estimated cover values. This characteristic is consistent with the general structure of peat bog vegetation, but the data obtained do not allow conclusions to be drawn regarding productivity, peat accumulation, or hydrological dynamics. The relationship observed between canopy closure and moss layer cover should be regarded as a descriptive observation and a hypothesis for future research.

Gaina-Lucina stood out through the presence of *Betula nana*, a taxon of high conservation value and relict character in the flora of Romania. Its presence is one of the main elements distinguishing the plot investigated and underlines the conservation interest of the site. However, a single relevé does not allow the size, structure, or trend of the population to be assessed across the entire peat bog, which is why this species requires distinct and repeated monitoring.

The 2026 phenological observations revealed seasonal successions of the main phenophases for most of the taxa monitored. The particularity observed in *Vaccinium myrtillus* at Gaina-Lucina, together with the differences recorded between the 2023 and 2026 observations, indicate the possibility of inter-annual variability, but do not allow the identification of a multi-year trend or of causal factors within the available dataset.

Overall, the study provides a descriptive floristic and phenological characterisation of the four plots investigated and highlights both the common elements of peat bog vegetation and the floristic particularity of Gaina-Lucina. The interpretation of the results is limited by the use of a single 25 m^2^ plot for each peat bog, by phenological monitoring over a single season, by the identification of some taxa only to genus level, and by the lack of direct hydrological and pedochemical measurements.

Repeating relevés at the same plots and expanding sampling to additional plots would allow a more robust characterisation of floristic variability. Determining *Sphagnum* species and integrating measurements of the hydrological regime, substrate characteristics, and light conditions would contribute to a better understanding of the factors associated with vegetation structure. For *Betula nana*, repeated monitoring of the Gaina-Lucina population represents a priority direction for research and conservation.

## Acknowledgments

The author thanks his wife, Elena, for her support and for her company during field trips, and for the help provided with some of the measurements.

